# Structural implications of glycosylation on the voltage-gated sodium channel β3-subunit

**DOI:** 10.1101/2025.02.27.640513

**Authors:** Christopher A. Beaudoin, Jerrard Hayes, Hengrui Liu, Sivakumar Namadurai, Michael J. Deery, Ethan A. August, Samuel Krasner, Samir W. Hamaia, Samantha C. Salvage, Christopher L.-H. Huang, Antony P. Jackson

**Author notes:** Corresponding author: APJ, Department of Biochemistry, Hopkins Building, Downing Site, Tennis Ct Rd, Cambridge CB2 1QW, United Kingdom. Authors contributed equally (Christopher A. Beaudoin, Jerrard Hayes, Hengrui Liu, Sivakumar Namadurai).

## Abstract

Voltage-gated sodium (Na_V_) channel α-subunits are modulated by associated β-subunits that affect their localization, trafficking and gating behaviour. The β-subunits are members of the immunoglobulin (Ig) domain family of cell-adhesion molecules and the interactions between their extracellular Ig-domains may modify channel clustering. The full-length β3-subunit can form cis trimers on the plasma membrane. The atomic resolution structure of a deglycosylated trimeric β3-subunit Ig-domain has been solved by X-ray crystallography. However, it is not clear whether this particular trimeric Ig-domain structure is plausible for cell-expressed, glycosylated β3-subunits. Here we use glycan profiling to confirm an extensive and heterogeneous pattern of β3-subunit glycosylation, with the majority of glycans being bi- and tri-antennary structures with one or two terminal sialic acids. Two tryptic peptides of the β3 Ig-domain are predicted to contain potential N-linked glycosylation sites. When the isolated, glycosylated full-length β3-subunit was trypsin-digested and analysed by LC-MS/MS, only one of these peptides - containing an N-linked glycosylation site at residue N95 and located close to the trimer interface - was identified in its unmodified form, suggesting that residue N95 is under-glycosylated. All-atom molecular dynamics simulations of the glycosylated, membrane-bound full-length β3 trimer confirmed that glycans can be accommodated with the Ig-domain trimer and indeed, may contribute to protein-membrane and inter-protomer interactions within the full-length, membrane-embedded trimer. Further biochemical studies are warranted to explore the interactions between oligomeric β-subunits with corresponding α-subunit sodium channels.

## 1. Introduction

Voltage-gated sodium channels (Na_V_1.1-1.9) initiate action potentials along nerve and muscle fibers(1). The Na_V_ channels comprise an ion-selective α-subunit, regulated by associated β-subunits, of which there are four distinct isoforms (β1-4)(2–4). The β-subunits are characterized by an N-terminal immunoglobulin (Ig) domain, a flexible linker region, a transmembrane domain, and a C-terminal intracellular tail (Figure 1A, B)(5). The β-subunits are structurally related to members of the Ig family of cell-adhesion molecules and have been implicated in both cis and trans cell-cell adhesion(6). Homophilic cis interactions may allow attachment between neighbouring channels, while homophilic trans interactions may permit contacts with cells across the extracellular matrix. Such cell adhesion may be important for cell migration, pathfinding, fasciculation and stabilising cell-cell contacts(7).

**Figure 1.**
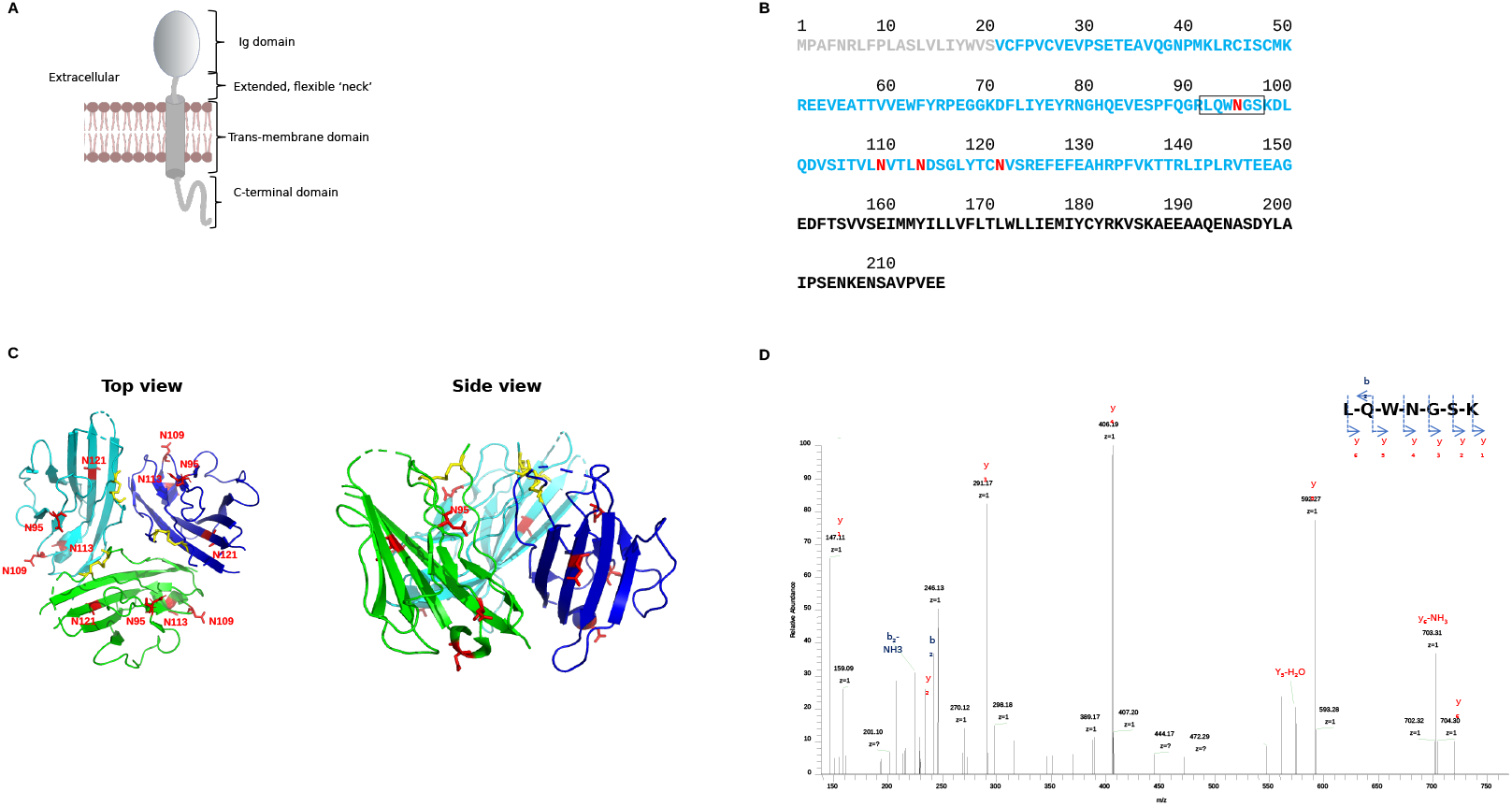
The β3-subunit. (A) Cartoon showing the key structural features of the β3-subunit. (B) The primary sequence of the β3-subunit. The ER targeting signal, removed in the endoplasmic reticulum, and not included in the MD simulations is coloured grey. The mature Ig domain sequence is shown in cyan. As discussed in the text, the four potential N-linked glycosylation sites highlighted in red. The tryptic peptide 92-98, containing potential N-linked glycosylation site N95 and discussed in the text is boxed. (C) The top and side view of the trimeric β3 Ig domain crystal structure (PDB: 4L1D). Individual trimers are separately coloured for clarity. Putative N-linked glycosylation sites are colored red and indicated. (D) MS/MS spectrum for doubly charged ion tryptic peptide 92-98 (LQWNGSK). There is a full set of C-terminal (y) fragment ions and the asparagine residue is deamidated.

Multiple biochemical and structural techniques reveal that the β3-subunit and its isolated Ig-domain can form trimers in cis, independently of Na_V_ α-subunits(8), and cell adhesion assays have ruled out homophilic trans interactions(9–11). Trimerization of β3 may also enhance spatially localized clustering of α-subunits on the plasma membrane(10). However, since little is known about the β3 interactions, further work may delineate the stability and biochemical roles of oligomerization of the β3 in cis homophilic interactions. As determined by x-ray crystallography, the β3 Ig-domain trimer interface is stabilised by hydrophobic interactions between the first six N-terminal residues of the mature protein. Notably, three valines (V25, V27, and V29) at the N-terminus form a hydrophobic network with the other two protomers(8). However, the β3 Ig-domain contains four potential N-linked glycosylation sites fitting the consensus sequence: Nx[S or T], where x can be any amino acid except proline (Figure 1B, C)(8, 12). The purified β3 Ig-domain runs as multiple bands on SDS PAGE, with apparent Mwts significantly higher than expected from the amino acid sequence alone and the migration anomaly is removed following PNGase treatment(8, 13). This indicates extensive and heterogeneous glycosylation on the β3 Ig-domain at some or all of the four sites. But the crystal structure of the trimeric β3-subunit Ig-domain was determined using a deglycosylated preparation. The question therefore arises whether the crystallographic trimer structure can accommodate these glycosylation patterns. If not, then its relevance to β3 cis oligomerisation is called into question.

Herein, high-performance liquid chromatography techniques were utilized to identify the major glycan species present on the β3 Ig-domain. Interestingly, mass spectrometry revealed that one of the four N-linked glycosylation sites, which is near the oligomerization interface, is not glycosylated. Using structural modelling and all-atom molecular dynamics simulations, we confirm that the presence of glycan trees is structurally compatible with the β3 Ig-domain trimer structure previously identified by X ray crystallography and that site-specific N-linked glycosylation may regulate interactions with Na_V_ α- and β-subunits. Further structural and functional implications for β3 in Na_V_ channel assembly are discussed.

## 2. Results

### 2.1. N-linked glycans on the β3 Ig-domain

As part of an ongoing study into β3-subunit expression in cancer cells, we transiently expressed the full-length β3-subunit with a C-terminal GFP-tag, in the C6 glioblastoma cell-line. LC-MS/MS experiments were performed on the immunoprecipitated β3 protein in separate duplicate experiments. Trypsin cleavage sites were predicted and matched with the returned spectra, and all peptide ends corresponded to predicted cleavage sites, confirming complete trypsin cleavage.

There are 9 tryptic peptides of length ≥ 4 amino acids within the β3 Ig-domain of which 8 were consistently detected with high confidence in two separate biological replicate experiments. One of the potential N-linked glycosylation sites (N95), occurs within tryptic peptide 92-98. The remaining three potential N-linked glycosylation sites (N109, N113, and N121), occur within tryptic peptide residue 98-124 (Figure 1B). Due to their anomalous and heterogeneous m/z values, glycosylated peptides are not easily identified by the Mascot scoring algorithm. In this context, it is interesting that peptide 98-124 was never detected in any of our MS/MS experiments, despite otherwise high overall peptide identification. By contrast, peptide 98-124 was consistently detected, and clearly indicated the presence of an non-glycosylated N95 residue (Figure 1D). Taken together with the SDS PAGE evidence noted above, this data suggests that the majority of the glycans are present on one or more of the three N-linked glycosylation sites present in peptide 98-124. Within the β3 Ig-domain trimer structure, residues N109, N113 and N121 are all prominently located on the Ig-domain surface, far from the trimer interface, where they would be particularly accessible to glycosylation enzymes. It is interesting to note that, whilst residue N95 is similarly surface-exposed in the monomeric Ig-domain, in the β3 Ig-domain crystal trimer structure, N95 occurs closer to the putative trimer interface (Figure 1C). Thus, uniquely for the trimer, N95 may be relatively hidden to the glycosylation enzymes during assembly in the endoplasmic reticulum.

### 2.2. Diverse glycan structures cover the β3 Ig-domain

To better understand the composition of N-linked glycans on the surface of β3, the β3 Ig-domain was expressed in HEK293 cells, purified from media supernatant, and HILIC HPLC, WAX analysis and exoglycosidase digestions of the Ig-like domain N-glycans were performed. Glycan analysis was performed by immobilization of β3 Ig-domain in SDS-PAGE gels and in-gel PNGase F release of N-glycans followed by fluorescent labeling with 2-aminobenzamide and HILIC UPLC analysis. Triplicate analysis of the labelled glycans showed that the results obtained were reproducible with minor differences in the relative abundances. As shown in Figure 2A, the majority of glycans, approximately 60-70%, contained one or two sialic acids with the majority of these being monosialylated. Human IgG1, which has well characterised glycan structures was used as a control glycoprotein to ensure that the glycan release and labelling occurred appropriately and bovine fetuin was used for comparison and determination of charged sialylated structures in WAX analysis. Sialidase treatment followed by WAX analysis was further used to analyse the charged content of β3 Ig-domain glycans and confirmed the presence of mainly neutral and mono-sialylated glycans, as the sialylated peaks migrated to the neutral fraction following treatment (see Figure 2B).

**Figure 2.**
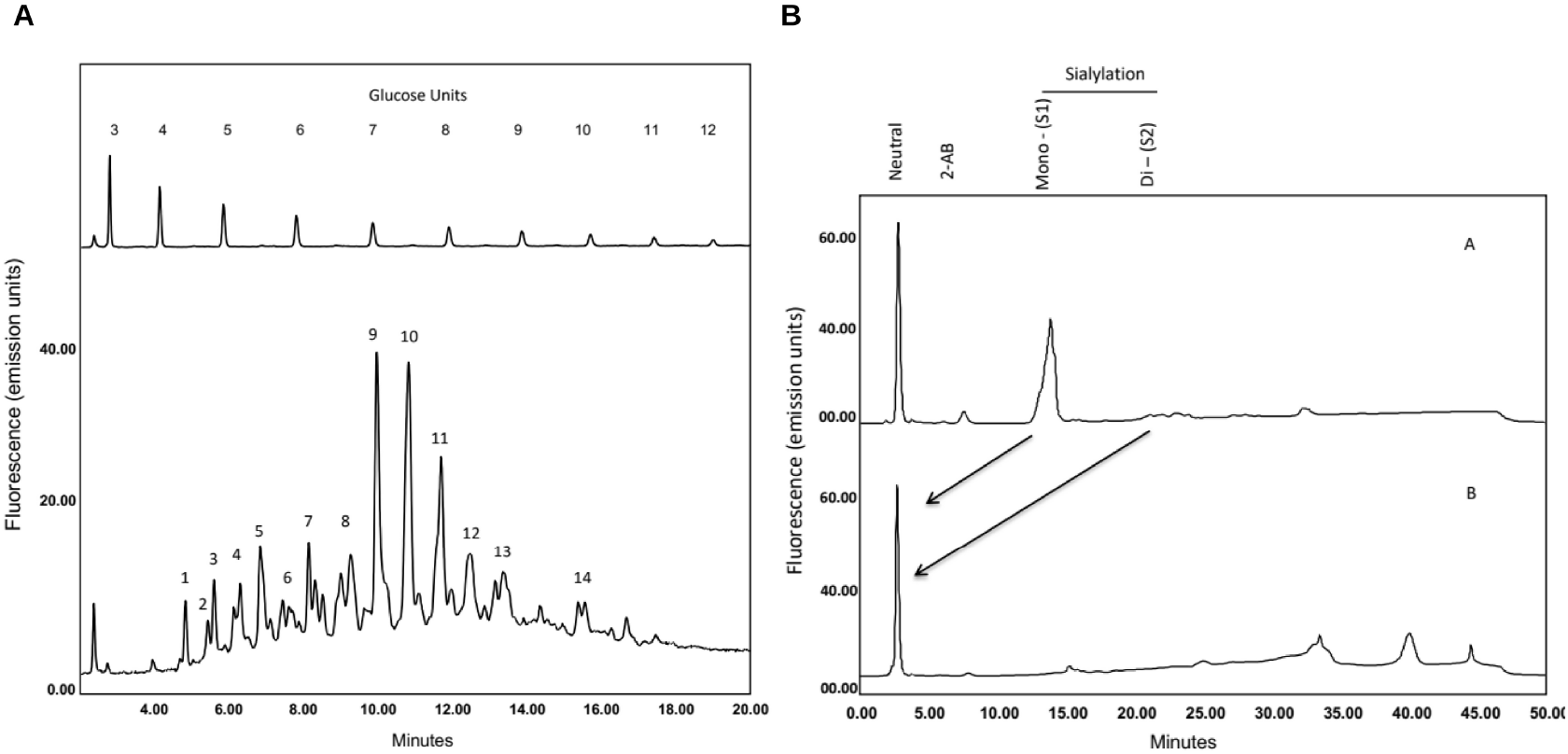
The β3 Ig-like domain displays complex and heterogenous glycosylation. (A) The major glycans were identified following enzymatic release and HILIC UPLC analysis and correspond to the structures shown in Table 1. GU values were assigned using internal 2-AB labelled dextran standards and integration using Waters Empower 3 software. Glycan structures and relative abundance are shown in Table I. Glycan release and analysis experiments were performed in triplicate. (B) WAX HPLC analysis of β3 Ig-like domain shows the glycans are predominantly neutral and mono-sialylated with minor amounts of disialylated structures. B. Sialidase treatment followed by WAX analysis revealed that peaks correspond to charged sialic acids. Bovine fetuin, which contains mono-, di-, tri- and tetra-sialylated glycans was used as standard to identify the sialylated species of β3 Ig-like domain (not shown).

To sequence the N-glycans of β3 Ig-domain and determine the monosaccharide compositions and linkage information a series of exoglycosidase digestions were performed and analysed by HILIC UPLC (see Figure 3). Panel digests including α-sialidase (ABS), β-galactosidase (BTG), α-fucosidase (BKF) and β-hexosaminidase (GUH) and α-mannosidase (JBM) which typically reduce N-glycans to common core mannose structures and this approach allowed the tentative identification of N-glycans (see Table 1). The glycans were primarily composed of complex bi- and tri-antennary structures, which are mainly core-fucosylated with varying degrees of sialylation and smaller amounts of oligomannose moieties (Figure 3). While glycosylation was found to be heterogenous, the most abundant glycans were identified as bi-antennary structures with one or two terminal sialic acids. See Table 1 for glycan structures identified and their relative abundance. These data provide novel insights into the organization of glycan moieties on the β3 surface and may be further used to infer the effect of glycosylation on the β3 structure and function.

**Table 1.**
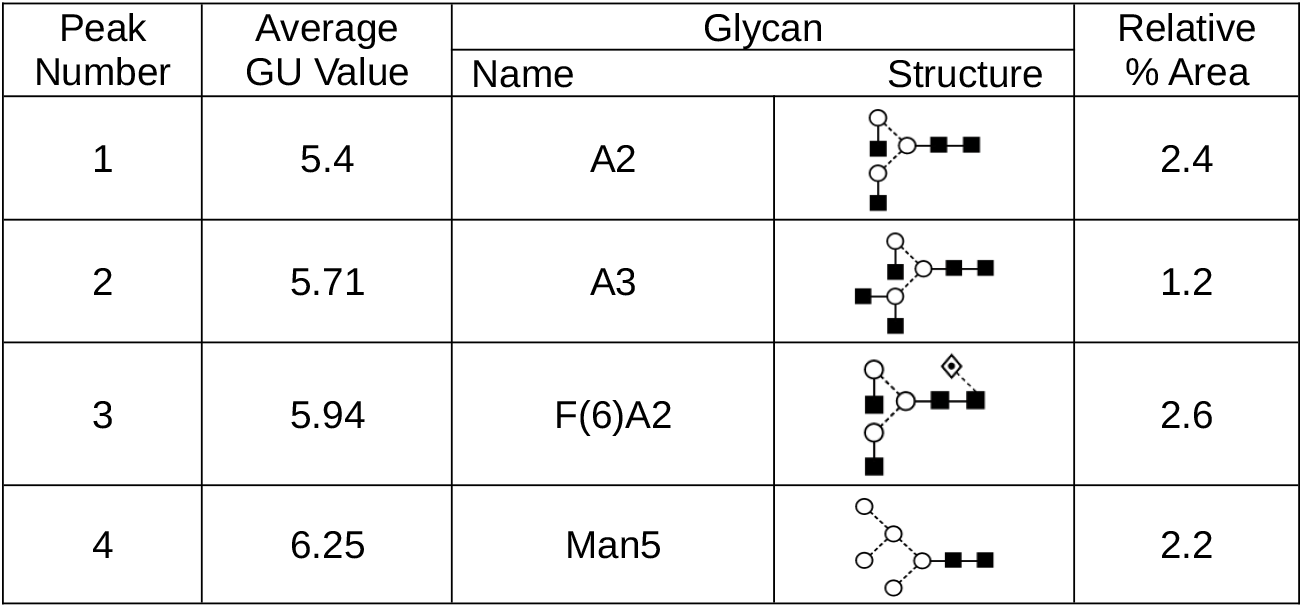
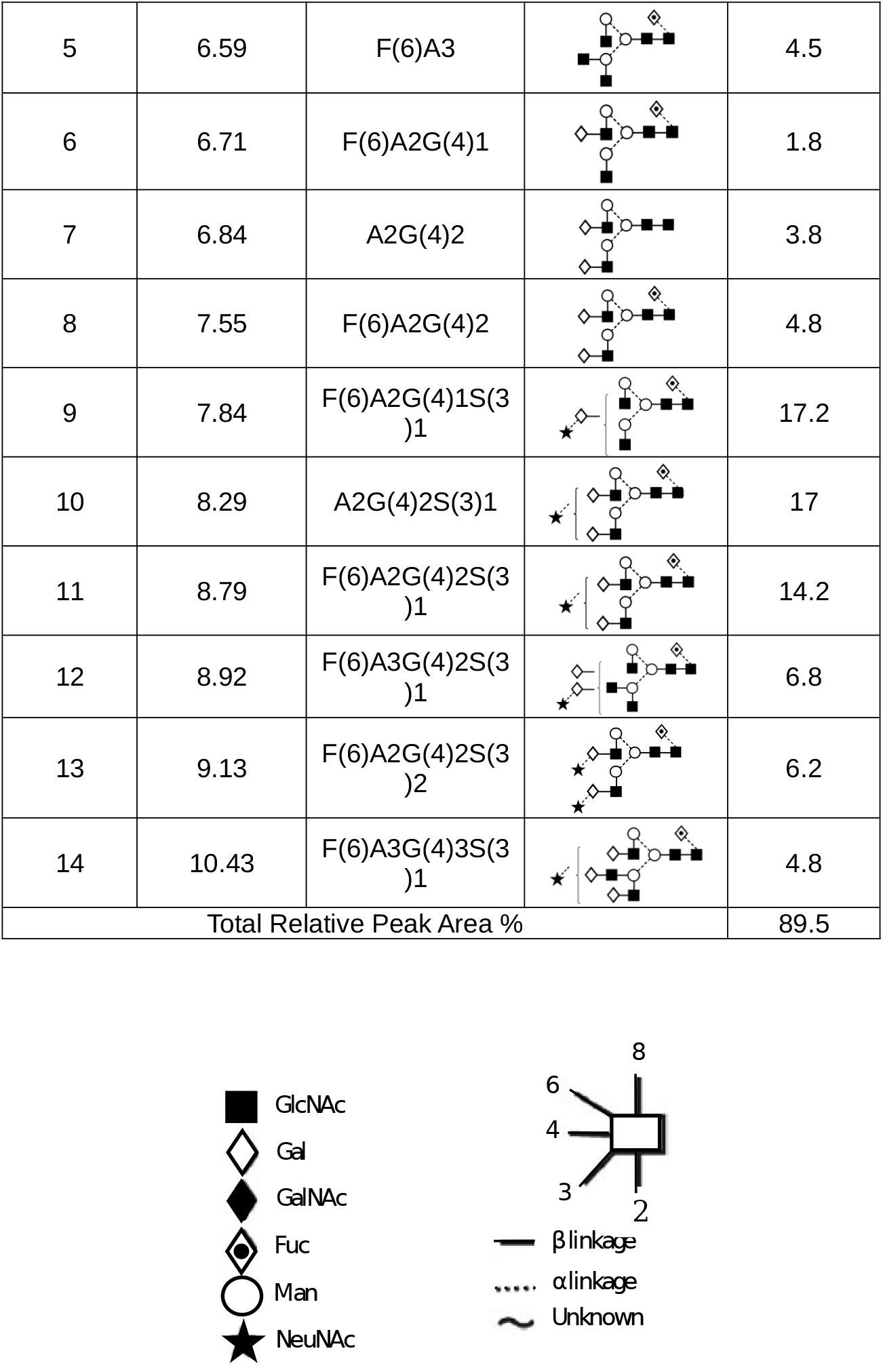
N-glycans of β3 Ig-like domain. Glycans were identified and quantified using a combination of HILIC UPLC, exoglycosidase digestions and WAX HPLC. Average GU (Glucose Unit) values and % areas are the mean of three individual glycan releases. Peak numbers correspond to peaks shown in Figure 2. Glycans are named according to the Oxford notation as illustrated in the legend below Table 1.

**Figure 3.**
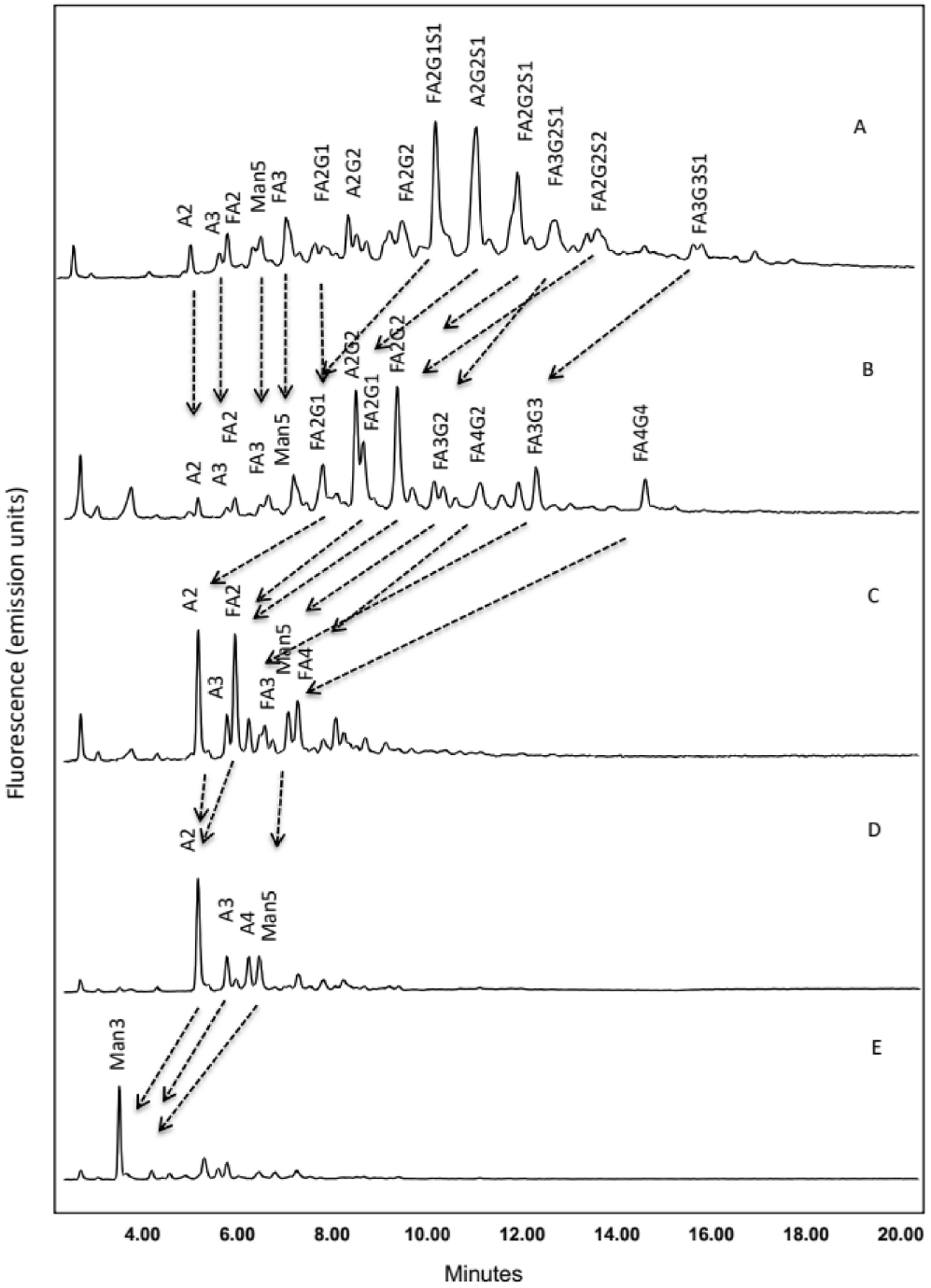
Exoglycosidase sequencing of the β3 Ig-like domain. Exoglycosidase sequencing of the β3 Ig-like domain N-glycans was used to identify the monosaccharide compositions and linkage information. Arrows indicate the migrations of peaks following exoglycosidase digestion and HILIC UPLC analysis A. Undigested β3 Ig-like domain glycan profile B. Arthrobacter ureafaciens sialidase (ABS) C. bovine testes-galactosidase (BTG) D. bovine kidney-fucosidase (BKF), E. jack bean β-N-acetylhexosaminidase (JBH). Peaks migrate to Man3 structures following digestion. Exoglycosidase sequencing was performed for each FcγR.

### 2.3. Shielding of β3 residues by N95 glycan

The conformational diversity of N-linked glycans on protein surfaces has been shown to affect protein-protein and other intermolecular interactions. The effective ‘cloud’ of density that is generated by glycans on protein surfaces has been utilized across all domains of life for various functions, such as preventing recognition by proteases or antibodies. Considering the relatively high density of N-linked glycans on the β3 surface and the proximity of the N95 glycan to the oligomerization interface, investigating the glycan density may shed light on oligomerization and interactions with the Na_V_ channel α-subunit. Therefore, the GlycoShield program was used to generate 30 distinct conformations of the N-linked glycans – using the most abundant glycan tree from the profiling in Section 2.2 as a template – on the β3 protomer Ig-domain surface using coarse-grained molecular dynamics simulations. The solvent accessible surface area (SASA) of each β3 residue during the simulations was recorded to determine the amount of ‘shielding’ across the protein surface. Both the fully and partially (without N95 glycan) glycosylated forms were analyzed to determine how the N95 glycan, specifically, may affect interactions with Na_V_ α- and other β3-subunits.

Initial structural models of the fully glycosylated β3 Ig-domain protomer (Figure 4A) and trimer (Figure 4B) did not reveal any apparent effect on oligomerization. However, after running GlycoShield and comparing SASA profiles between the fully and partially glycosylated forms, only a few residues were found to be differentially shielded (Figure 4C). Notably, glycosylation of the N95 residue was found to shield the N-terminal valines (V25, V27, and V29), which are the primary residues involved in trimerization, and other residues on the Ig-domain surface (Figure 4D). When aligning the fully and partially glycosylated forms into the β3 trimer crystal structure (Figures 4E and 4F), the conformations of the N95 glycan can be seen directly overlapping with the Ig-domain of another β3 protomer. Therefore, glycosylation of the N95 residue may interfere with oligomerization via two mechanisms: shielding of the oligomerization interface and deterring nearby β3 Ig-domains from interacting. The partially glycosylated β3 protomer may, thus, be more accommodating for trimerization.

**Figure 4.**
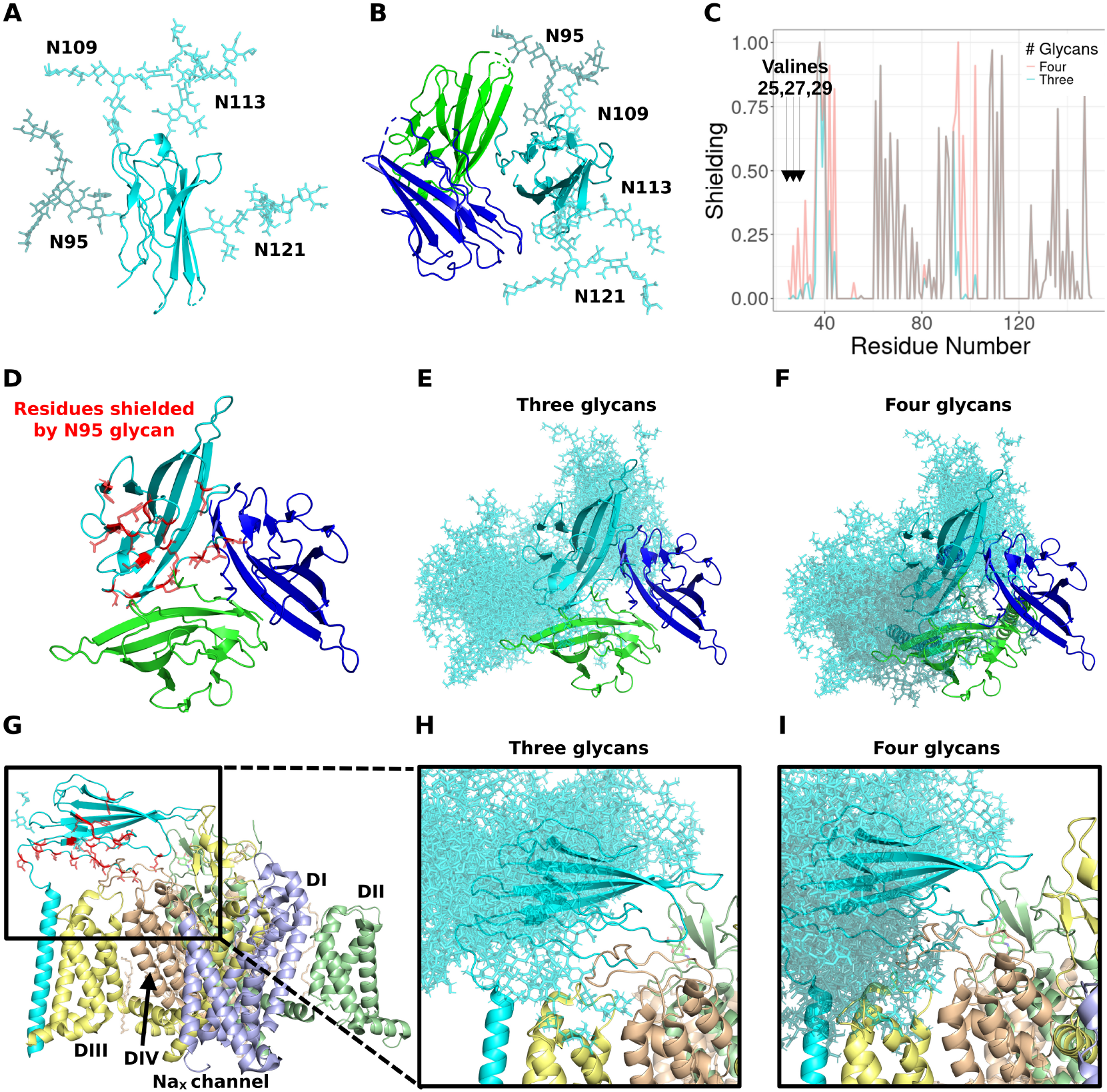
Effect of glycosylation on β3 inter-protomer interactions The β3 trimer is shown with each protomer colored in a different shade of blue and the glycans are shown in varying colors in a representative plasma membrane (grey) (A). Valines at the oligomeric interface are depicted as spheres (B). One pair of glycans at N109 on one protomer

The fully and partially glycosylated forms of β3 were also aligned to the cryo-EM structure of β3 bound to the Na_X_ channel α-subunit (PDB: 7TJ9). Similarly to the trimer crystal structure, conformations of the N95 glycan are found overlapping with key regions of the Na_X_ channel (Figure 4G). For example, the N95 glycan overlaps an extracellular loop on domain IV of the Na_X_ channel, which is both responsible for channel gating and binding to the β3 Ig-domain. Importantly, glycans may be identified on cryo-EM structures by the presence of extra electron density near NX[S,T] sites on a protein surface. Interestingly, in the Na_X_-β3 cryo-EM structure, N-linked glycans are found at all four NX[S,T] sites on the β3 surface. This structural information indicates that glycosylation of the N95 residue may occur after complexing between β3 and the α-subunit, since the shielding of the interaction interface may interfere with complexing to the α-subunit. However, additional sites on both β3 and the α-subunit may nevertheless interact and lead to complexing. Further analysis of the structural features of β3 glycosylation may shed light on its dynamics in protein-protein interactions.

#### Glycans modulate inter-protomer and protein-membrane interactions

Structural modelling of the interactions that occur between the β3 protomers, N-linked glycans, and cell membrane may further delineate the roles of N-linked glycosylation. Therefore, using an identical glycan tree from Section 2.3, models of fully (glycans at N95, N109, N113, and N121) and partially (no glycan at N95) glycosylated full-length β3 trimers in a representative bilayer membrane were generated. All-atom molecular dynamics simulations of the models were then conducted in triplicate for 500 ns each to better understand the intermolecular interactions and, more specifically, the structural effects of glycans on β3 oligomerization over time.

The glycans modelled at the N109, N113, and N121 residues were noted to point outwards from the trimer and either parallel to or towards the plasma membrane (Figure 5A), and the N95 glycan was found pointing directly away from the membrane to accommodate the close packing of protomers. In investigating the protein-protein interactions of the trimer models, all simulations supported the previously stated assertion that the N-terminal strands are the primary interfacing residues (Figure 5B)(8). Three valines – V25, V27, and V29 – were found to form a hydrophobic cluster, which is common among branched aliphatic side chain amino acids(14), at the interface and stabilized the association of the three protomers (Figure 5C). Although the V25 residues in the crystal structure were facing inwards towards the respective protomers and away from the interface, the simulations suggest that the terminal valines rotate to interact at the interface while being anchored to the protomer by the neighboring C26 residue involved in a disulfide bond with C48. When comparing the effect of glycosylation on the oligomerization interface using the distance between V29 residues as a metric, one of the simulations with four glycans was found to lead to dissociation of the trimer (Figure 5D). All other simulations retained an intact trimer interface throughout, although the inter-valine distance did vary slightly more between models with zero or four glycans than three glycans. These results suggest that three glycans may effectively stabilize inter-protomer interactions.

**Figure 5.**
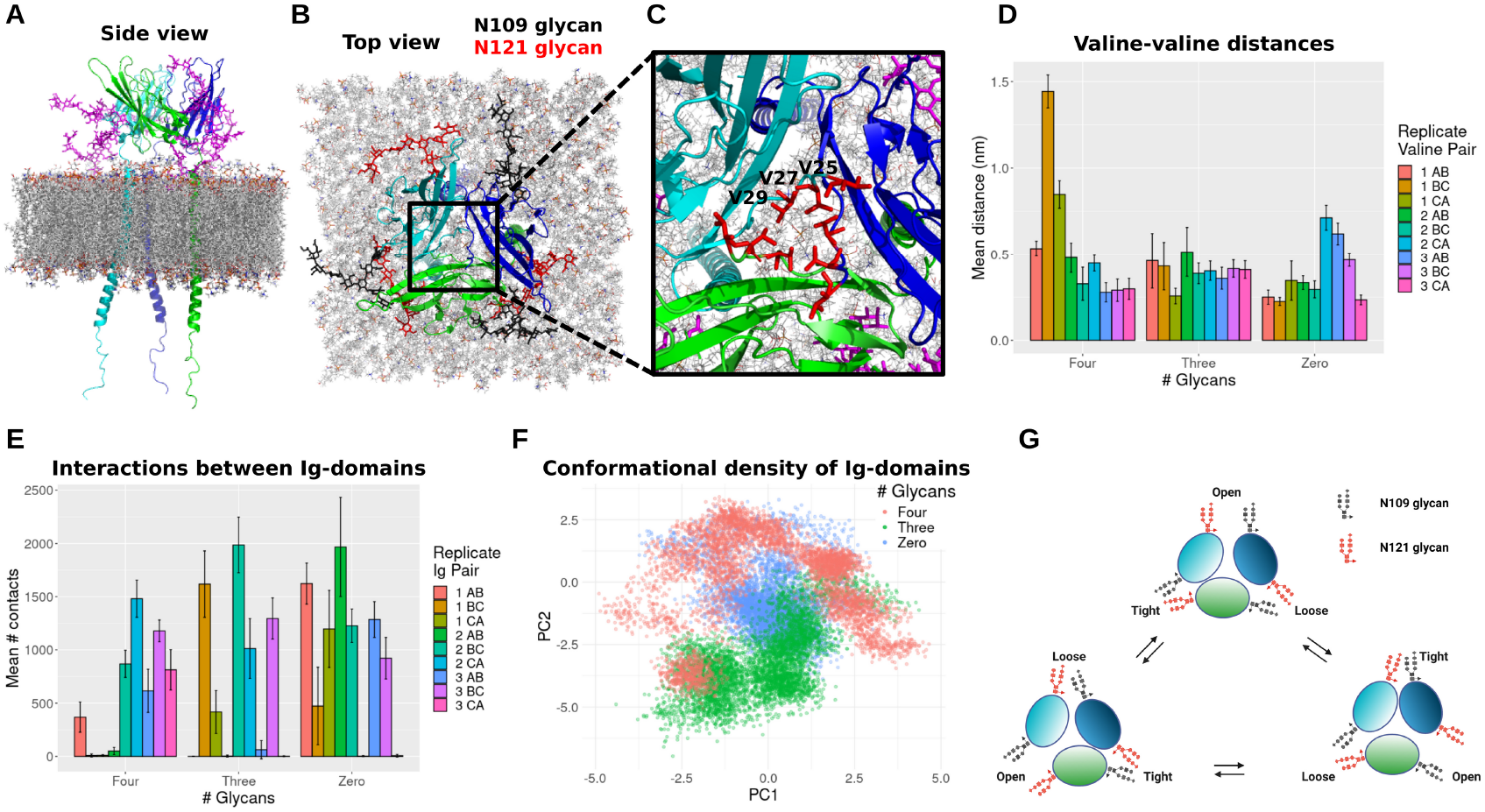
Effect of glycosylation on β3 inter-protomer interactions (A) The β3 trimer is shown with each protomer colored differently and the glycans are shown in magenta in a representative plasma membrane (grey). (B) One pair of glycans at N109 on one protomer and N121 on a neighboring protomer are shown in black and red, respectively. (C) Valines at the oligomeric interface are depicted as red stick. (D) Mean distances between the V29 residues over the last 50 ns of each simulation are shown as a bar plot. (E) The number of contacts (atoms < 6 Å from one another) between β3 protomer Ig-domains over the last 50 ns of each simulation are shown as a bar plot. (F) A principal component analysis (PCA) plot of conformations sampled throughout all replicate simulations is depicted with each glycan mode colored differently (G) A schematic portraying the differences in recorded symmetry between the glycosylated and non-glycosylated Ig-like domains is featured.

Interactions between the β-strands that comprise the Ig-domains (T33-F153) may also stabilize and contribute to trimerization. As shown in Figure 5E, most of the simulations revealed a bias towards interactions between only two of the three β3 protomers. Such data indicates that there is an asymmetry among β3 trimer Ig-Ig contacts. Notably, the modelled glycans at N109 on one protomer and N121 of the neighboring protomer were found to interact (Figure 5B). Calculation of the average radial distribution between the sugar moieties at these sites revealed step-wise differences between protomer sets (Figures 6A-B) - i.e. the glycans of two protomers were found to interact strongly, while another set of two was found to interact less, and the third set even less than the second. Interestingly, comparing the radial distributions between Ig-domains of the protomers revealed that the glycans were found to more consistently contribute to asymmetric inter-protomer distance for only the models without the glycan at N95 (Figures 6C-E). The glycans were found to interact most between the closest Ig-like domains, and the glycans with the second closest distance were found to interact second closest in Ig-like domains in two of the three simulations (Figures 6D). Interactions between the glycans for fully glycosylated (including at N95) models did not reflect the Ig-Ig interaction profiles (Figures 6B and 6E). The non-glycosylated Ig-Ig β-strand distances were shown to be more symmetrical in two of the three simulations (Figure 6C). However, one run was more asymmetric than all three glycosylated runs, suggesting that asymmetry may occur regardless of glycosylation states. The data above suggest that partial glycosylation (absence of the N95 glycan) may help stabilize asymmetry of the trimer more effectively than with four or zero glycans (Figure 5G). Therefore, inter-protomer interactions may be partially regulated by the selective presence of glycans.

**Figure 6.**
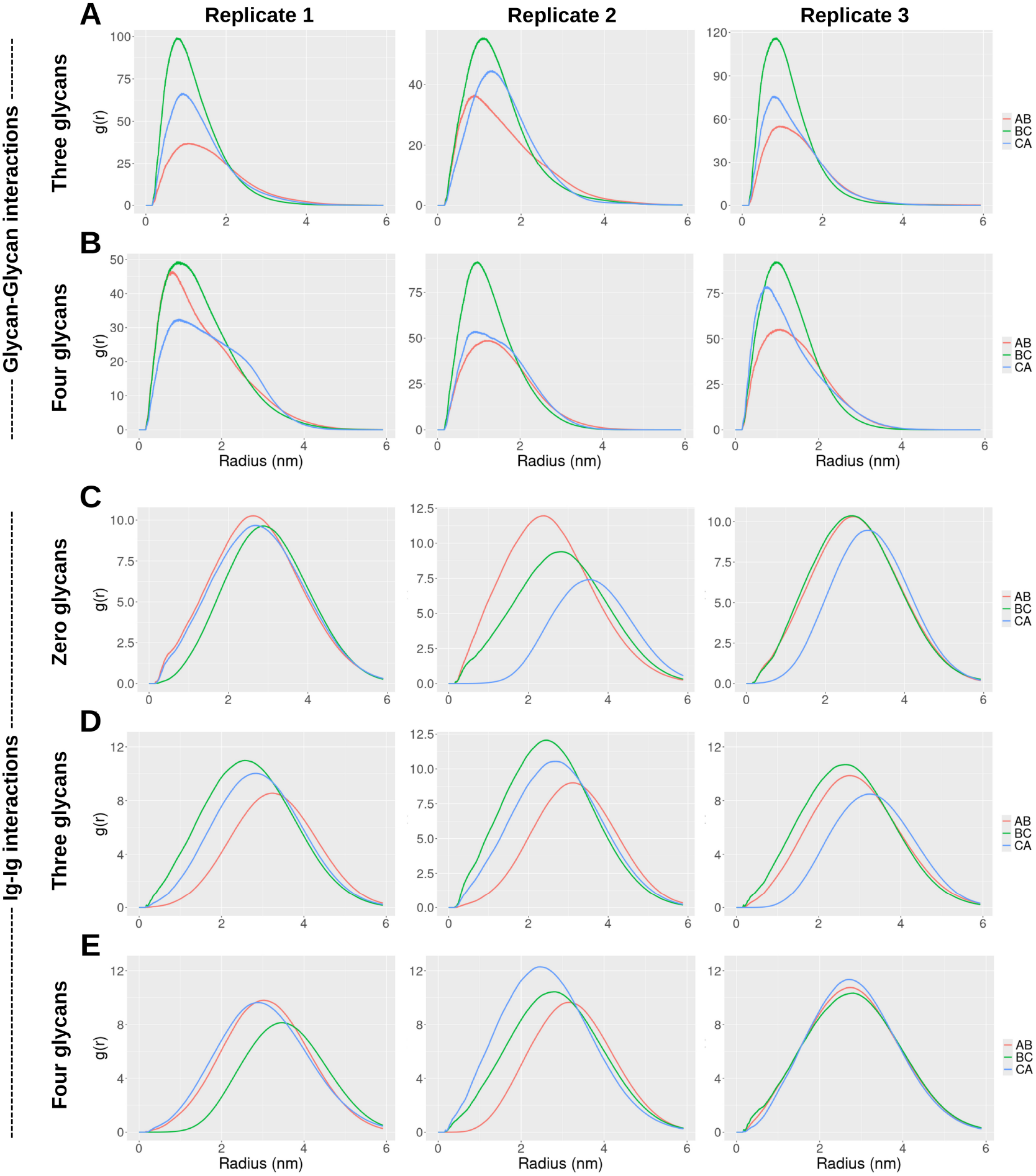
Inter-protomer interactions The radial distributions generated by three independent simulations for: (A-B) distances between the glycans at N109 and N121 and (C-E) between each pair of protomer Ig-domains.

Conformational clustering of the β3 trimer Ig-domains was additionally performed to determine the effect of glycans on complex stability. As shown in Figure 5F, a principal component analysis of the β3 Ig-domain conformational states revealed noticeable overlaps between the models with four, three, and zero glycans, suggesting multiple conformational spaces sampled by both glycosylated and non-glycosylated states. Despite the overlap, distinct regions in the PCA space are primarily occupied by each glycosylation state. The models with four glycans appear to explore a broader and slightly shifted conformational space, which may suggest increased structural variability. The zero glycan cluster forms a compact core, likely reflecting the base conformations sampled in the absence of glycosylation. Interestingly, the three glycan clusters are generally placed furthest from the four glycans clusters, which might indicate that these states, although both glycosylated, represent distinct conformational assemblies. Considering that no region is heavily populated with conformational states from all three simulation modes but each mode shares a region with both of the others, the progression from zero to three to four glycans might reflect unique and functional glycan-induced modulation of shared conformational states. Therefore, glycosylation of β3 protomers may alter the conformational landscape, potentially pointing to functional differences.

Inspection of the putative glycosylation at N113 revealed that the glycan trees are primarily involved in interactions with the plasma membrane. The glycans may act as ‘stilts’ upon which the protomers are shielded from interactions with the plasma membrane and can stabilize into the trimeric state (Figures 7A-B). Notably, the glycosylated Ig-domains were found at a further distance from the plasma membrane when compared to non-glycosylated forms – the three glycan models being the furthest (Figure 7C). These data suggest that the increased destabilization induced by the N95 glycan may lead to less emphasis on the N113 glycan interactions with the cell membrane. In addition to the glycans at the N113 sites, glycans at the N109 and N121 sites were closer to the membrane in the Ig-domains that were further from the membrane (Figures 7D-E). Therefore, all glycan sites may contribute to modulating the distance between the cell membrane and glycosylated Ig-domains. Considering the findings above, each β3 glycan likely regulates various aspects of Ig-Ig and Ig-membrane interactions.

**Figure 7.**
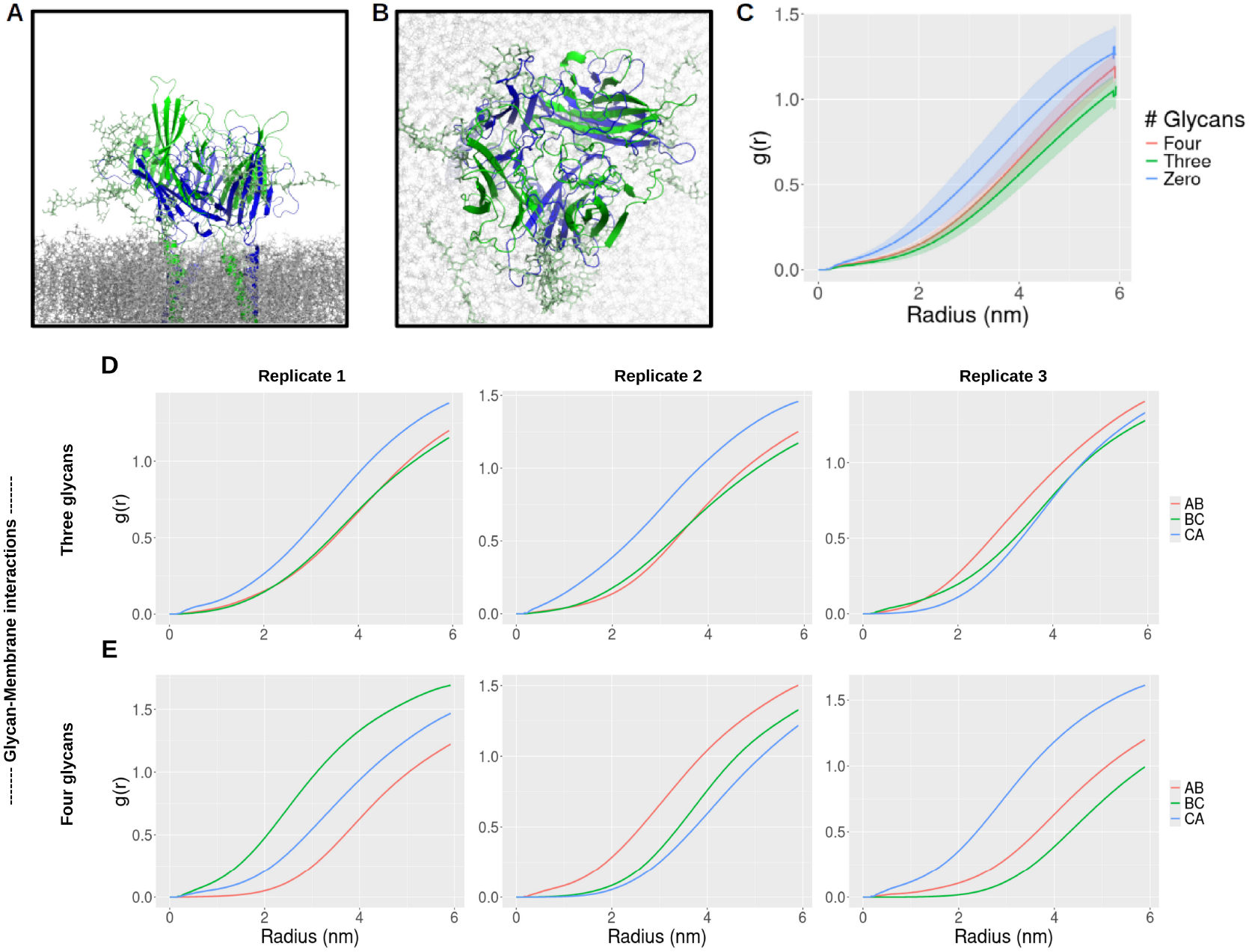
Interactions between the membrane, glycans, and β3 protomers The top (A) and side (B) view of representatives of both the partially glycosylated (green) and non-glycosylated (blue) β3 trimers at the end of the respective simulations are shown. (C) The mean radial distributions between the glycosylated and non-glycosylated Ig-like domains and the membrane lipids are shown shaded with the standard deviation. (D-E) Radial distributions between both glycans (N109 and N121) and membrane lipids for each simulation are shown.

## Discussion

Previous MD simulations on β3 monomers and oligomers predicted several cis oligomeric states, including those similar to the crystal structure formation, but also heterogeneous states in which the protomers were interacting via different interfaces. However, these simulations, did not consider glycosylation patterns(15). Our results confirm that a trimer, in which the Ig-domains interact as in the crystal structure can form even when glycosylated. Indeed, MD simulations of the glycosylated β3 trimer revealed that glycans at three sites (N109, N113, N121) may have a stabilizing effect on the trimer, while glycosylation at another site (N95) may destabilize the trimer. The glycan at N95 may either prevent trimerization via shielding of the oligomerization interface or sterically hinder the Ig-domains from associating. These findings corroborate with the mass spectrometry data, which indicate that the N95 residue is not consistently glycosylated. Glycosylation of the N109 residue on one protomer was predicted to interact with the putative glycan at the N121 residue on another protomer, thus potentially increasing contacts between neighbouring Ig-like domains. Furthermore, glycosylation at the N113 residue were found to stabilize the Ig-like domains standing away from the plasma membrane lipids, potentially acting as ‘stilts’ upon which protomers can adopt the configuration seen in the crystal structure, whilst simultaneously protecting the otherwise exposed ‘neck’, connecting the Ig and TM domains against proteolytic attack. Glycan analysis of the recombinant β3 Ig-domain revealed glycosylation to be heterogenous. However, the major glycans identified were biantennary structures with mainly one terminal sialic acid and smaller amounts with two sialic acids. Approximately 90% of the total glycans were tentatively identified. While it is likely that these glycan structures play a role in structure and stability of the Na_V_ channel it is also possible the large amount of these negatively charged glycan structures play a role in ion channel selectively and function.

When bound to the NaV channel α-subunit, the β3-subunit usually induces a depolarising shift in steady-state voltage dependence of inactivation and may enhance the rate of recovery from inactivation(10, 16). There are currently no detailed atomic-resolution structures for β3 in association with classical Na_V_ channel α-subunits. Recently however, the cryo-EM structure of β3-subunit associated with the non-classical sodium channel Na_X_ has been published(17). In this structure, the β3-subunit Ig-domain binds to the extracellular loop regions of the α-subunit domain III voltage-sensing domain (VSD) via the same N-terminal regions as forms the β3 trimer interface. If the β3-subunit binds to other Na_V_ channel α-subunits in the same way, then a given β3-subunit would not be able to simultaneously bind an α-subunit and form a homophilic trimer. However, we have previously noted that the heart-specific sodium channel, Na_V_1.5, contains a unique N-linked glycosylation site, the glycosylation of which would prevent the binding of the β3 Ig-domain and force it to lie outward, away from the domain III VSD(18). In this case, the β3 Ig-domains may then be able to form trimers and facilitate larger cis-interacting Na_V_1.5 clusters. An analogous situation may also apply to the β2 and β4-subunits, whose Ig-domains contain a free cysteine residue that forms a disulfide bond with most Na_V_ channel α-subunits. But the β2 and β4-subunits Ig-domains can also form cis-mediated disulfide bonded, homophilic dimers using the same cysteine residue. Thus, for most Na_V_ channels, β2 and β4 can either dimerise homophilically or bind the α-subunit but cannot simultaneously do both. Remarkably, however, Na_V_1.5 lacks the necessary cysteine residue to partner β2 or β4. Hence, we have suggested that these unusual isoform-specific features of Na_V_1.5, acting together, facilitate cis clustering via the β2, β3 or β4 subunits, each in their distinct β-isoform specific manners. Such clusters are likely to be particularly important in cardiomyocytes, for example on the opposing perinexal membranes of the intercalated disks(6). It will therefore be interesting to now explore the structural feasibility of these ideas by further computational and experimental investigations.

## Materials and Methods

### β3 culture transfection, purification, and LC-MS/MS preparation

The preparation of the β3 Ig-like domain for the glycan profiling was performed as previously described(8). The protein expression of full-length human β3, in order to determine protein-protein interactions, was conducted using C6 cells as part of a separate series of experiments. The C6 cell line was provided by Dr. Sarah McGuire from Prof. Kevin Brindle’s lab at Cancer Research UK Cambridge Institute. C6 cells were cultured in DMEM medium (Invitrogen) supplemented with 10% FBS at 37 °C and 5% CO2. The expression vector plasmids of β3-GFP (green fluorescence protein) or GFP were constructed by cloning. The expression vector with GFP was transfected into cells at 0.5 mg/ml using polyethylenimine (PEI, 1µg/1mL). The observation of GFP in cells and western blotting of the GFP was used to determine transfection efficiency.

The cells were lysed with 1% Triton X-100, 0.6 M Tris buffer, pH 7.4, 0.88 % NaCl, and cOmplete™ protease inhibitor. Anti-GFP beads immunoprecipitation was used to purify the protein using ChromoTek GFP-Trap® Agarose. A bead slurry of 25 µL was transferred into a 1.5 mL tube and equilibrated with 500 uL lysis buffer 3 times. Lysate of 500 µL was mixed with beads and rotated at 4 °C for 1 h. Then the beads were washed with PBS 3 times. The binding proteins were eluted by 50 µL acidic glycine elution buffer (0.2 M glycine pH 2.5) twice. The eluted samples were neutralized by adding 10 µL neutralization buffer (1 M Tris pH 10.4). The neutralized eluted samples were processed for the subsequent mass spectrometry experiment. The sample processing and mass spectrometry as well as the analysis of the raw data were conducted by the Cambridge Centre for Proteomics. Immunoprecipitated proteins of β3-GFP or GFP were further processed for the proteomic-mass spectrum. The samples were run in a NuPAGE Bis-Tris gel, and the bands were transferred into an Eppendorf tube. The bands were cut into 1 mm^2^ pieces, destained, reduced (DTT) and alkylated (iodoacetamide) and subjected to enzymatic digestion with trypsin overnight at 37 °C. After digestion, the supernatant was pipetted into a sample vial and loaded onto an autosampler for automated LC-MS/MS analysis.

### Liquid Chromatography Tandem Mass Spectrometry (LC-MS/MS)

Duplicate LC-MS/MS experiments were performed using a Dionex Ultimate 3000 RSLC nanoUPLC (Thermo Fisher Scientific Inc, Waltham, MA, USA) system and a Q Exactive Orbitrap mass spectrometer (Thermo Fisher Scientific Inc, Waltham, MA, USA). Separation of peptides was performed by reverse-phase chromatography at a flow rate of 300 nL/min and a Thermo Scientific reverse-phase nano Easy-spray column (Thermo Scientific PepMap C18, 2 μm particle size, 100A pore size, 75 μm i.d. x 50 cm length). Peptides were loaded onto a pre-column (Thermo Scientific PepMap 100 C18, 5 μm particle size, 100A pore size, 300 μm i.d. x 5mm length) from the Ultimate 3000 autosampler with 0.1% formic acid for 3 minutes at a flow rate of 15 μL/min. After this period, the column valve was switched to allow the elution of peptides from the pre-column onto the analytical column. Solvent A was water + 0.1% formic acid and solvent B was 80% acetonitrile, 20% water + 0.1% formic acid. The linear gradient employed was 2-40% B in 40 minutes. Further washing and equilibration steps gave a total run time of 60 minutes.

The LC eluant was sprayed into the mass spectrometer by means of an Easy-Spray source (Thermo Fisher Scientific Inc.). All m/z values of eluting ions were measured in an Orbitrap mass analyzer, set at a resolution of 35000 and scanned between m/z 380-1500. Data-dependent scans (Top 20) were employed to automatically isolate and generate fragment ions by higher energy collisional dissociation (HCD, NCE:26%) in the HCD collision cell and measurement of the resulting fragment ions was performed in the orbitrap analyser, set at a resolution of 17500. Singly charged ions and ions with unassigned charge states were excluded from being selected for MS/MS and a dynamic exclusion window of 20 seconds was employed. Text files generated from the raw data were converted to mgf files, and the files were used in the sequencing mapping to the SCN3B gene (UniProt: Q9NY72)(19) sequence. Trypsin cleavage sites were predicted using the ExPASy PeptideCutter(20) with the trypsin-based statistical model(21), and only sites predicted with confidence scores of 100% were selected(22).

### N-linked Glycan Release and 2-aminobenzamide (2-AB) Labelling of HEK293 expressed β3 Ig domain

Recombinant β3 Ig domain expressed in HEK293 cells was analysed by SDS PAGE and the N-linked glycans were released and labelled by the SDS-PAGE immobilisation and release method described by Royle *et al*.(23). Approximately 100 μg of individual receptors were resolved by SDS PAGE and visualised by Coomassie® blue staining. Bands were excised from SDS PAGE gels and glycans were released by incubation at 37 °C overnight with 0.1 mU recombinant peptide-N-glycosidase F (Prozyme). Released glycans were extracted from gel slices and labeled with 2-aminobenzamide (2AB; Ludger) according to the manufacturer’s instructions and purified using normal phase resin columns. (Phytips, Phynexus).

### Hydrophilic Interaction Ultra Performance Liquid Chromatography (HILIC UPLC)

2-AB labelled glycans were prepared as 80 % (v/v) acetonitrile solutions for analysis by HILIC UPLC using a Waters BEH-Glycan 1.7 μm (150 mm x 2.1 mm) column and fluorescence detection (excitation at 420 nm and emission at 330 nm) as previously described ref. All separations were performed using a Waters Acquity H-Class UPLC instrument over a 30 min period at 40 °C using 50 mM ammonium formate pH 4.4 as solvent A and 100 % (v/v) acetonitrile as solvent B. 2-AB-labelled dextran was used as an internal calibration standard and for the generation of glucose unit (GU) values. Retention times for 2-AB labelled dextran peaks were used to fit a fifth order polynomial distribution curve using Waters Empower 3 software, which provided the means for converting chromatographic retention times into standardized glucose unit (GU) values as previously described. Glycans were named according to the Oxford notation.

### Weak Anion Exchange HPLC (WAX HPLC)

WAX HPLC was used to analyse the extent of terminal sialylation of β3 Ig domain N-glycans and for fractionation of charged structures. A Waters Biosuite® DEAE 10μm AXC (7.5 mm x 75 mm) column was used for separation of charged carbohydrate structures. Solvent A was 100 mM ammonium acetate pH 7 in 20 % (v/v) methanol and solvent B was 20 % (v/v) acetonitrile. All WAX HPLC separations were performed on a Waters Alliance 2695 instrument with online Waters 4795 fluorescence detector. Carbohydrate samples were separated over a 30 min gradient. The relative proportions of terminal charged glycan structures were determined by comparison to N-linked glycans released from bovine fetuin, which contains neutral, mono-, di-, tri-, and tetra-sialylated structures. For WAX-fractionation experiments glycan pools were separated by WAX HPLC and peaks from this analysis were manually collected. Glycans were dried, reconstituted in MilliQ water and analysed by HILIC UPLC. This 2-dimensional approach was used to deconvolute the complex N-glycan pools.

### Exoglycosidase Panel Digests

Exoglycosidase arrays were used to sequence the N-Glycans of β3 Ig domain. Using this technique, sequence, composition and linkage specificity information was obtained. Fluorescently labelled glycans were routinely digested in 50 mM sodium acetate, pH 5.5 at 37 °C overnight in a volume of 10 μL using a selected panel of exoglycosidase enzymes. The following day enzymes were removed using 10 kDa spin filters (Pall Corp, NY, USA). Digested 2-AB-labelled glycans were analysed by HILIC UPLC as described previously. The specificities of the exoglycosidase enzymes used are well-established and specific monosaccharides were removed as follows: terminal sialic acid in all linkages: α(2,6), α(2,3), α(2,8) is removed with 1 mU/μL Arthrobacter ureafaciens sialidase (ABS)(Prozyme); terminal galactose monosaccharides were removed using 0.5 mU/μL bovine testes β-galactosidase (BTG)(Prozyme), which releases both (1,3)- and (1,4)-linked galactose; terminal N-acetylglucosamine (GlcNAc) monosaccharides in β(1,4) linkage were released with 40 mU/μl Streptococcus pneumoniae hexosaminidase (GUH)(Prozyme); core (1,6)-fucose was selectively removed using 1 mU/μL bovine kidney-fucosidase (BKF)(Prozyme); terminal non-reducing end fucose in α(1-3)- and α(1-4)-linkage was removed using 0.004 mU/μL almond meal-fucosidase (AMF)(Prozyme) and βGalNAc and βGlcNAc were removed using 50 mU/μL jack bean β-N-acetylhexosaminidase (JBH)(Prozyme).

### Full-length β3 trimer modelling

The full-length AlphaFold2 β3 protomer structure model (UniProt: Q9NY72)(19, 24, 25) was aligned to each protomer of the experimentally-determined β3 structure model (PDB: 4L1D)(8). Signal peptide residues (M1-P24) were removed. Foldit Standalone(26) was then utilized to energetically minimize the main and side chain energies and orient the protomer flexible linker and transmembrane domains parallel to one another to be conducive for placement in a representative plasma membrane. Two successive runs of FoldX RepairPDB and one run of FoldX Optimize were conducted on the finalized trimeric model to fix torsion angles, Van der Waals clashes, and minimize total energy(27). The protonation states of β3 trimer residues were validated with H++ at a pH of 7 using the TIP3P water model and, otherwise, default settings(28). The most prevalent glycan tree structure was then modelled onto each of the three determined sites per protomer using the CHARMM-GUI Glycan Reader and Modeller(29). The glycosylated model was then placed in a representative mammalian cell membrane, based on the work by Ingólfsson et al.(30) - using PPM 2.0(31) and the CHARMM-GUI Membrane Builder(32). The lipid bilayer membrane is composed of DSM (upper leaf: 21.0%, inner leaf: 10.0%), POPC (35.0%, 15.0%), DOPC (3.5%, 1.5%), POPE (5.0%, 20.0%), DOPE (2.0%, 5.0%), POPS (0%, 15.0%), POPI (0%, 5.0%), POPA (2.2%, 0%), and CHOL (31.3%, 28.5%)(18). The E176 on each protomer was protonated to study its effects based on previous reports.

### Molecular dynamics simulations

All-atom molecular dynamics simulations were prepared and conducted using GROMACS 2021.3(33) installed with the University of Cambridge High Performance Computing resources. Periodic boundary conditions were established in a cubic box set at 115×115×115 Å. All systems were solvated in 150 mM NaCl at a zero net charge based on the TIP3P model(34) and were conducted at 310 K(35). Long-range electrostatic interactions were calculated using the Particle-mesh Ewald method, and Coulomb interactions and van der Waals interactions cut-offs were set to 12 Å(36). The LINCS algorithm was used to constrain molecular bonds(37). Following steepest descent minimization, all systems were subjected to six series of a 125 picosecond NPT equilibration ensemble with temperature coupling using velocity rescaling(38) and pressure coupling using the Parrinello-Rahman method(39). All simulations were run using the CHARMM36 force field (C36 FF) with 2 femtosecond time steps(40, 41). Triplicate productions of 500 nanoseconds were conducted for the non-glycosylated, partially glycosylated, and fully glycosylated membrane-bound full-length β3 trimers.

Trajectories were visualized and snapshot structures were extracted with VMD(42). The 50 nanoseconds before convergence of the simulations of the runs were excluded in downstream analyses to account for the influence of equilibration of the systems(43). The RMSD, RMSF, radial distribution functions, and atom-atom distances were extracted from the simulation trajectories using the gmx rms, gmx rmsf, gmx rdf, and gmx dist (with contact prediction defined as distance threshold < 6 Å) GROMACS modules, respectively(44). The data were plotted using R version 3.6.3.

## Acknowledgments

We thank the University of Cambridge High Performance Computing Team for their advice and assistance.

## Data availability

All data is available upon request.

## Author contributions

CAB: Conceptualization, Methodology, Validation, Formal analysis, Investigation, Writing - Original Draft, Writing - Review & Editing. JH: Conceptualization, Methodology, Validation, Formal analysis, Investigation, Writing - Original Draft, Writing - Review & Editing. SN: Conceptualization, Methodology, Validation, Formal analysis, Investigation. HL: Conceptualization, Methodology, Validation, Formal analysis, Investigation, Writing - Original Draft, Writing - Review & Editing. MD: Methodology, Validation, Formal analysis, Investigation. EAA: Methodology, Validation, Formal analysis. SK: Methodology, Validation, Formal analysis. SWH: Investigation, Writing - Original Draft, Writing - Review & Editing. SCS: Conceptualization, Investigation, Writing - Original Draft, Writing - Review & Editing, Funding acquisition. CLHH: Conceptualization, Supervision, Project administration, Funding acquisition. APJ: Conceptualization, Methodology, Supervision, Project administration, Funding acquisition.

